# ALSgeneScanner: a pipeline for the analysis and interpretation of DNA NGS data of ALS patients

**DOI:** 10.1101/378158

**Authors:** Alfredo Iacoangeli, Ahmad Al Khleifat, William Sproviero, Aleksey Shatunov, Ashley R Jones, Sarah Opie-Martin, Ersilia Naselli, Isabella Fogh, Angela Hodges, Richard J Dobson, Stephen J Newhouse, Ammar Al-Chalabi

## Abstract

Amyotrophic lateral sclerosis (ALS, MND) is a neurodegenerative disease of upper and lower motor neurons resulting in death from neuromuscular respiratory failure, typically within two years of first symptoms. Genetic factors are an important cause of ALS, with variants in more than 25 genes having strong evidence, and weaker evidence available for variants in more than 120 genes. With the increasing availability of Next-Generation sequencing data, non-specialists, including health care professionals and patients, are obtaining their genomic information without a corresponding ability to analyse and interpret it. Furthermore, the relevance of novel or existing variants in ALS genes is not always apparent. Here we present ALSgeneScanner, a tool that is easy to install and use, able to provide an automatic, detailed, annotated report, on a list of ALS genes from whole genome sequence data in a few hours and whole exome sequence data in about one hour on a readily available mid-range computer. This will be of value to non-specialists and aid in the interpretation of the relevance of novel and existing variants identified in DNA sequencing data.

## Introduction

Amyotrophic lateral sclerosis (ALS) is a progressive neurodegenerative disease, typically leading to death within 2 or 3 years of first symptoms. Many gene variants have been identified that drive the degeneration of motor neurons in ALS, increase susceptibility to the disease or influence the rate of progression [1]. The ALSoD webserver [2] lists more than 120 genes and loci which have been associated with ALS, although only a subset of these have been convincingly shown to be ALS-associated [3], demonstrating one of the challenges of dealing with genetic data–interpretation of findings. Next-generation sequencing provides the ability to sequence extended genomic regions or a whole-genome relatively cheaply and rapidly, making it a powerful technique to uncover the genetic architecture of ALS [4]. However, there remain significant challenges, including interpreting and prioritizing the found variants [5] and setting up the appropriate analysis pipeline to cover the necessary spectrum of genetic factors, which includes expansions, repeats, insertions/deletions (indels), structural variants and point mutations. For those outside the immediate field of ALS genetics, a group that includes researchers, hospital staff, general practitioners, and increasingly, patients who have paid to have their genome sequenced privately, the interpretation of findings is particularly challenging.

The problem is exemplified by records of *SOD1* gene variants in ALS. More than 180 ALS-associated variants have been reported in *SOD1* [2]. In most cases, the basis of stating the variant is related to ALS is simply that it is rare and found in *SOD1*.

Neither of these conditions is sufficient for such a statement to be made. The p.D91A variant, for example, reaches polymorphic frequency in parts of Scandinavia, and yet has been convincingly shown to be causative of ALS. A few variants have been modelled in transgenic mice, shown to segregate with disease or have other strong evidence to support their involvement [6–7] but most do not have such support. Rare variation can be expected to occur by chance, and its existence in a gene is not evidence of relationship to a disease, making interpretation of sequencing findings difficult. Although various tools are available to predict the pathogenicity of a protein-changing variant, they do not always agree, further compounding the problem.

We therefore developed ALSgeneScanner, an ALS-specific framework for the automated analysis and interpretation of DNA sequencing data. The tool is specifically targeted for use by people with knowledge outside genetics.

## Methods

ALSgeneScanner is part of the DNAscan suite [8]. Figure 1 shows the pipeline main steps. A detailed description and benchmark of its analysis components has been previously published [8]. ALSgeneScanner uses, among others, Hisat2 [9] and BWA-mem [10] to align the sequencing data to a reference genome, Freebayes [11] and GATK Haplotype Caller [12] to call SNVs and small indels, Manta [13] and ExpansionHunter [14] for the detection of large structural variants (bigger than 50bps) and repeat expansions.

**Figure 1.**
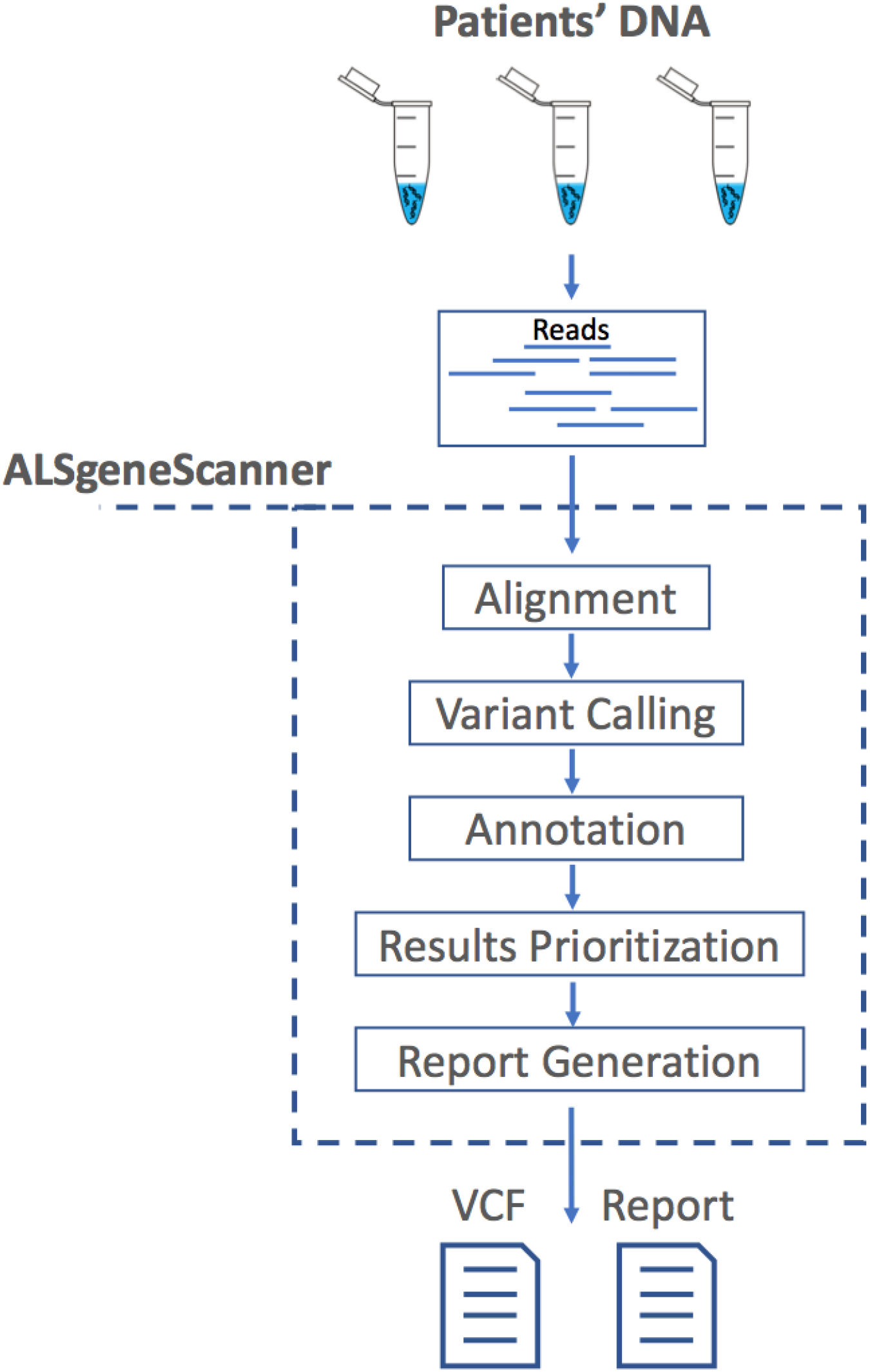
ALSgeneScanner pipeline main steps. From sequencing data in fastq format to the report generation of the results.

### Gene and loci prioritization

ALSgeneScanner groups genes and loci associated with ALS into three classes: i) genes for which a manual scientific literature review identified a strong and significant association with the disease or influence on the phenotype in ALS (see Table 1), ii) genes in which variants of clinical significance have been reported on ClinVar [15] and for which no contradictory interpretation is present, and iii) genes for which any association evidence has been submitted to ALSoD [2]. The union of these three sets of genes is used to restrict the genome analysis. However, ALSgeneScanner allows the user to use a custom list of genes.

**Table 1:** List of ALS genes identified by literature review

| Gene | Associated ND | Phenotype influence | Key Reference (doi) |
| --- | --- | --- | --- |
| **ANG** | ALS/PD |  | 10.1038/srep41996. |
| **ANXA11** | ALS |  | 10.1126/scitranslmed.aad9157 |
| **APOE** |  | Longer survival | 10.1212/WNL.58.7.1112 |
| **ATXN2** | ALS |  | 10.1038/nature09320 |
| **CAMTA1** |  | Shorter survival | 10.1001/jamaneurol.2016.1114 |
| **C21orf2** | ALS |  | 10.1038/ng.3622 |
| **C9ORF72** | FTD/ALS | Primarily bulbar onset | 10.1016/j.neuron.2011.09.011. |
| **CCNF** | FTD/ALS |  | 10.1038/ncomms11253 |
| **CHCHD10** | FTD/ALS |  | 10.1093/brain/awu138 |
| **DAO** | ALS |  | 10.1073/pnas.0914128107 |
| **DCTN1** | ALS |  | 10.1212/01.WNL.0000134608.83927.B |
| **EPHA4** |  | Longer survival | 10.1038/nm.2901 |
| **FIG4** | ALS |  | 10.1038/ejhg.2016.186 |
| **FUS** | FTD/ALS | Early age of onset and shorter survival | 10.1126/science.1165942 |
| **HNRNPA1** | ALS |  | 10.1038/nature11922 |
| **IDE** |  | Shorter survival | 10.1001/jamaneurol.2016.1114 |
| **MATR3** | ALS |  | 10.1038/nn.3688 |
| **MOBP** | ALS |  | 10.1038/ng.3622 |
| **NEK1** | ALS |  | 10.1038/ng.3626 |
| **OPTN** | ALS |  | 10.1126/science.aaa3650 |
| **PFN1** | ALS | Limb onset | 10.1038/nature11280 |
| **PGRN** | FTD/ALS |  | 10.1126/science.1077209 |
| **SARM1** | ALS |  | 10.1038/ng.3622 |
| **SCFD1** | ALS |  | 10.1038/ng.3622 |
| **SOD1** | ALS | Limb onset, early age of onset and shorter survival | 10.1038/362059a0 |
| **SPG11** | ALS |  | 10.1126/science.aaa3650 |
| **SQSTM1** | FTD/ALS |  | 10.1001/archneurol.2011.250 |
| **SETX** | ALS |  | 10.1086/421054 |
| **TAF15** | ALS |  | 10.1073/pnas.1109434108 |
| **TARDBP** | FTD/ALS |  | 10.1126/science.1154584 |
| **TBK1** | ALS |  | 10.1038/ng.3622 |
| **TUBA4A** | FTD/ALS |  | 10.1016/j.neuron.2014.09.027 |
| **UBQLN2** | FTD/ALS |  | 10.1038/nature10353 |
| **UNC13A** | ALS | Shorter survival | 10.1038/ng.3622 |
| **VAPB** | ALS |  | 10.1086/425287 |
| **VCP** | FTD/ALS |  | 10.1016/j.neuron.2010.11.036 |
| **8p23.2** | ALS |  | 10.1038/ng.3622 |
| **1p34-rs3011225** |  | Late age of onset | 10.1016/j.neurobiolaging.2012.07.017 |

### Variant prioritization

The pathogenicity prediction programmes, SIFT, Polyphen2 HDIV, Polyphen2 HVAR, LRT, MutationTaster, MutationAssessor, Fathmm, PROVEAN, Fathmm-MKL coding, MetaSVM and CADD are used to prioritize variants. A variant is scored X where X is equal to the number of tools which predict it to be pathogenic. A higher priority is given to variants which are reported to be “likely pathogenic” or “pathogenic” on ClinVar. For each tool we used the authors’ recommendations for the categorical interpretation of the variants. For each variant, the score ranges between 0 and 11 according to the number of computational tools (11 in total) that predict it to be pathogenic.

### Whole-genome sequencing

The whole-genome sequencing (WGS) sample used to assess the computational performance of ALSgeneScanner was sequenced as part of Project MinE [16]. Venous blood was drawn from patients and controls and genomic DNA was isolated using standard methods. DNA integrity was assessed using gel electrophoresis. All samples were sequenced using Illumina’s FastTrack services (San Diego, CA, USA) on the Illumina Hiseq X platform. Sequencing was 150bp paired-end performed using PCR-free library preparation, and yielded ~40x coverage across each sample.

### Whole-exome sequencing

To assess the computational performance of ALSgeneScanner we used the Illumina Genome Analyzer II whole exome sequencing of NA12878 (ftp://ftp-trace.ncbi.nih.gov/1000genomes/ftp/technical/working/20101201_cg_NA12878/NA12878.ga2.exome.maq.raw.bam)

### VariBench and ClinVar datasets

To assess our variant prioritization approach, we used a set of non-synonymous variants from the VariBench dataset [17] for which the effect is known and all ALS-associated non-synonymous variants stored in ClinVar. The VariBench variants are not ALS genes specifically, but because they are all annotated depending on whether or not they are deleterious, the general principles of the method could be tested. The dataset includes VariBench protein tolerance dataset 1 (http://structure.bmc.lu.se/VariBench/tolerance_dataset1.php) comprising 23,683 human non synonymous coding neutral SNPs and 19,335 pathogenic missense mutations [18].

### Evaluation of performance

Receiver Operating Characteristic (ROC) curves and their corresponding area under the curve statistic (AUC) were calculated using *easyROC* [19]. Accuracy, Precision and Sensitivity are defined as in Equation 1 where *T_p_* is true positives, *F_p_* false positives, *F_n_* false negatives and *T_n_* true negatives.

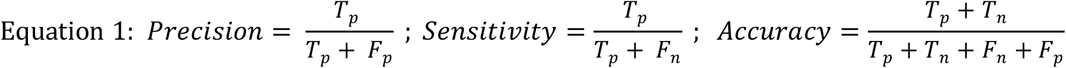

### Hardware

All tests were performed on a single, mid-range, commercial computer with 16GB RAM and an Intel i7-670 processor.

### Output

Resulting variants are reported in a tab delimited format to favour practical use of worksheet software such as iWork Number, Microsoft Excel or Google Spreadsheets.

### Software

ALSgeneScanner is available on GitHub [20] (https://github.com/KHP-Informatics/DNAscan). The repository provides detailed instructions for tool usage and installation. A bash script for an automated installation of the required dependencies is also provided as well as Docker [21] and Singularity [22] images for a fast and reliable deployment. A Google spreadsheet with the complete list of genes and loci used by ALSgeneScanner is publically available to visualize and comment (see Github repository).

## Results

Manual literature review identified 36 genes and 2 loci (Table 1) with strong and reproducible supporting evidence of association with ALS or influence on phenotype. ClinVar reported SNVs and small indels in 44 genes and 7 structural variants ranging in size from 3 to 50 million base pairs. ALSoD reported variants in 126 genes and loci. The union of these sets of genes contained 150 genes and loci. The Venn diagram in Figure 2 shows the overlap between the three sets.

Using a midrange commercial computer (4 CPUs and 16 gigabytes of RAM), (Figure 3) ALSgeneScanner could analyse 40x WGS data of one individual in about 7 hours using 12.8 Gb of RAM, and whole-exome sequencing data in 1 hour and 20 minutes using 8.5 Gb of RAM.

**Figure 2.**
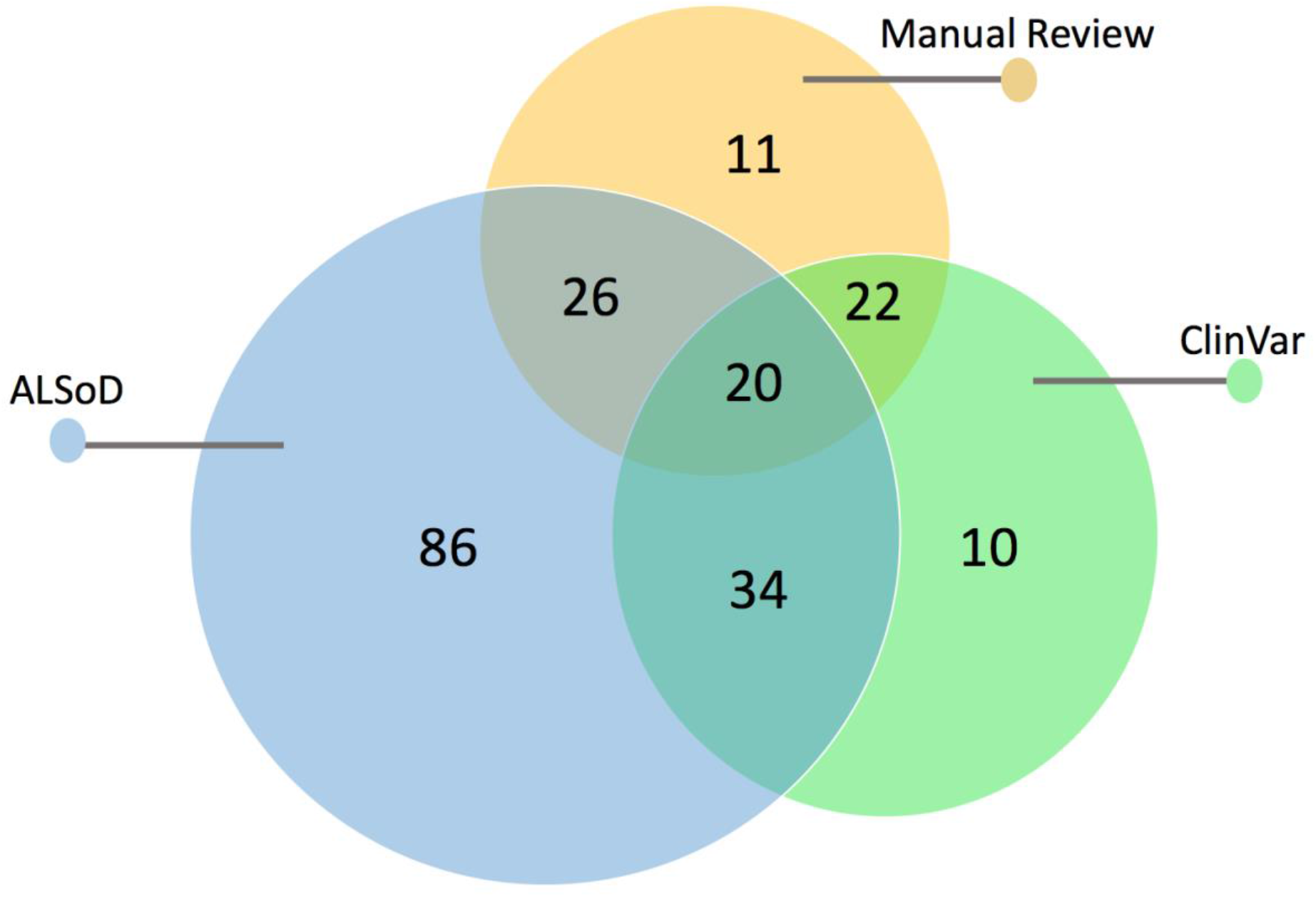
Venn diagram of the ALS related genes that we selected in our literature review, found in the ALSoD webserver and in the ClinVar database.

**Figure 3.**
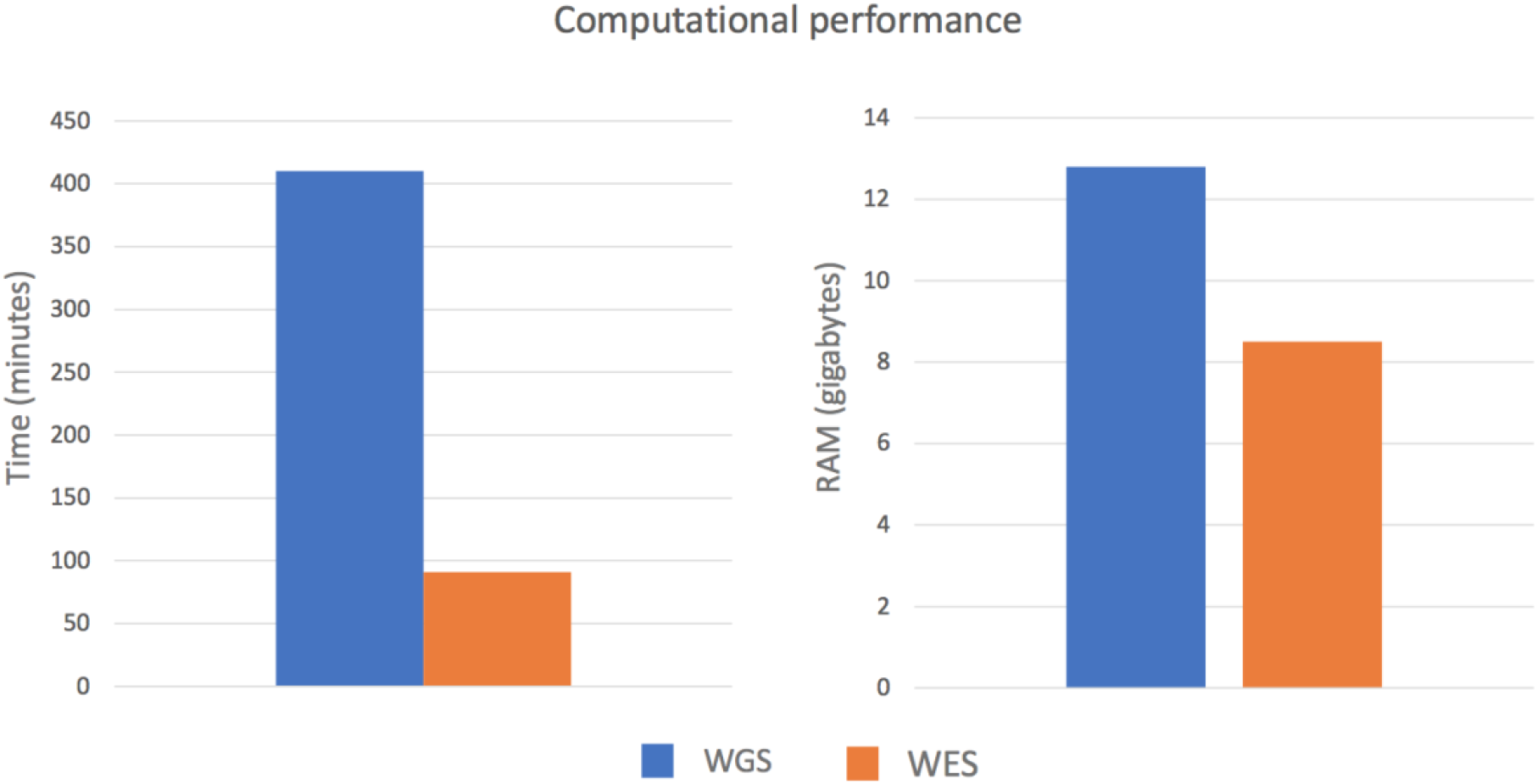
Computational performance of the pipeline to process whole genome sequence and whole exome sequence data from fastq file to the generation of the final result report.

We tested the computational score that the tool used to rank variants on two datasets. The VariBench dataset which includes ~40,000 variants of known effect (pathogenic and benign), and on the ALS associated ClinVar entries. Figure 4 shows the results on the two datasets and Table 2 precision, sensitivity and accuracy of the method in function of the chosen threshold for the VariBench variants. The ROC curve for the VariBench dataset (Figure 4, AUC = 0.90) suggests a cut-off equal to 9 which maximises the accuracy (0.827) however, a lower or higher cut-off can be chosen to reach a better precision or sensitivity according to the user’s needs. For example, for diagnostics a higher sensitivity is generally required and a cut-off equal to 5 would increase the sensitivity to 0.90 (Table 2). The ROC curve for the ClinVar variants suggests a cut-off equal to 7. Comparing to the VariBench variants, ClinVar ALS variants are more difficult to assess. The AUC for such variants is 0.82 (Figure 4) and the accuracy for the ideal cut-off is 0.75 (Table 2). This performance drop can be explained by the following factors: first the uncertainty in the ClinVar entries. ClinVar provides the community with an infrastructure to allow researchers to store their clinical observations, but the quality checks are very limited and the only filter we have adopted in this study to select the variants was the absence of contradictory entries. The second is the difficulty that available computational tools have in assessing the effect of ALS related variants [3, 23], in part because the mechanism of ALS is unknown, and in part because at least some of the variants result in a toxic gain of function that is difficult to understand or model.

**Figure 4.**
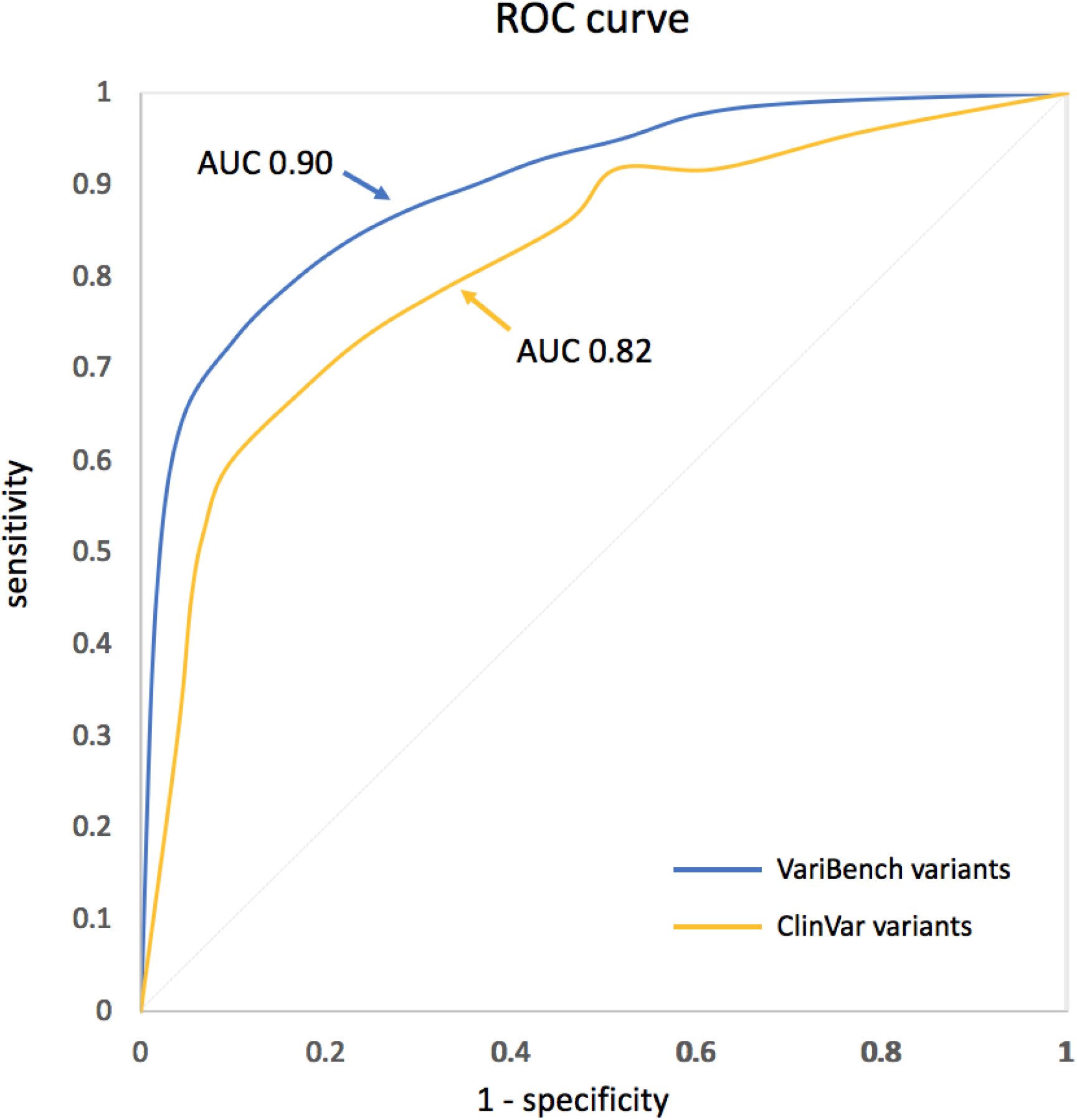
ROC curve of the performance of ALSgeneScanner on the two datasets.

**Table 2:** ALSgeneScanner variant prioritization performance

| Score | VariBench variants |  |  | ClinVar ALS variants |  |  |
| --- | --- | --- | --- | --- | --- | --- |
|  | Precision | Sensitivity | Accuracy | Precision | Sensitivity | Accuracy |
| 0 | 0.430 | 1 | 0.430 | 0.612 | 1 | 0.613 |
| 1 | 0.507 | 0.990 | 0.581 | 0.659 | 0.957 | 0.670 |
| 2 | 0.549 | 0.978 | 0.644 | 0.707 | 0.949 | 0.728 |
| 3 | 0.580 | 0.950 | 0.682 | 0.745 | 0.949 | 0.770 |
| 4 | 0.618 | 0.928 | 0.721 | 0.754 | 0.889 | 0.754 |
| 5 | 0.653 | 0.900 | 0.751 | 0.798 | 0.812 | 0.759 |
| 6 | 0.692 | 0.875 | 0.779 | 0.832 | 0.761 | 0.759 |
| 7 | 0.736 | 0.841 | 0.801 | 0.863 | 0.701 | 0.749 |
| 8 | 0.783 | 0.796 | 0.817 | 0.911 | 0.615 | 0.728 |
| 9 | 0.845 | 0.731 | 0.827 | 0.926 | 0.538 | 0.691 |
| 10 | 0.919 | 0.635 | 0.819 | 0.931 | 0.462 | 0.649 |
| 11 | 0.954 | 0.436 | 0.748 | 0.925 | 0.316 | 0.565 |

## Discussion

We have developed ALSgeneScanner, a fast, efficient and complete pipeline for the analysis and interpretation of DNA sequencing data in ALS, targeted for use by non-geneticists. The method is able to distinguish pathogenic from non-pathogenic variants with 83% accuracy and reports findings in a simple format, able to be exported for further analysis. With the decreasing costs and increasing availability of next generation sequencing, health care professionals and motivated patients are progressively more likely to have whole genome sequence data available, without the tools to interpret findings. An automated system to provide a meaningful report therefore has a potentially important part to play in giving patients ownership of their data and arming them with the knowledge to understand it.

Omictools [24], a web database where available bioinformatics tools are listed and reviewed, lists over 7000 such tools for next generation sequencing, including more than 100 pipelines; given the great interest in this field, new tools are frequently released. As a result, designing a bioinformatics pipeline for the analysis of next generation sequencing data, keeping the system simple to use on a standard computer and translating the output into a format that is easily understood, is not trivial, and requires specialised expertise. The computational effort and the informatics skills required to use typical pipelines can dramatically limit the use of next generation sequencing data. Adequate high-performance computing facilities and staff specialised in informatics are not always present in medical and research centres. Furthermore, the use of cloud computing facilities, which could theoretically provide unlimited resources, is not always possible due to privacy and ownership issues, cost and the expertise required for their use. To this end, ALSgeneScanner is computationally light as it can run on a midrange commercial computer, is easy to use since it performs sophisticated analyses using few command lines (see Box 1) and is comprehensive, including the necessary analyses to identify all known ALS associated genetic factors. Finally, a tab-delimited output, in which the analysis results are enriched with information from several widely used databases such as ClinVar and ALSoD, as well as information from our manual literature review, and the graphical visualization utilities integrated in the pipeline as part of DNAscan [8], favour an easily accessible interpretation of the results.

#### Box 1: Fast deployment and basic usage instructions for Linux based systems

##### Deployment

After downloading DNAscan^*^ (https://github.com/KHP-Informatics/DNAscan) and Annovar (http://www.openbioinformatics.org/annovar/annovar_download_form.php), you can deploy ALSgeneScanner by running one simple script that you can find in/path/to/DNAscan/scripts:

~~~
$./install_ALSgeneScanner.sh /path/to/DNAscan /path/to/annovar $num_cpus
~~~

Where $num_cpus is the number of cpus available on the machine, e.g. 4. This script will download and install all dependencies as well as create the necessary index files. The latter might require a little time depending on the number of CPUs used. To find out the number of CPUs available on your computer use the command nproc.

##### Usage

To run the whole ALSgeneScanner pipeline on paired-end reads in fastq format stored in data1.fq.gz and data2.fq.gz, you can run the following command running the main DNAscan scrip in /path/to/DNAscan/scripts :

~~~
$./DNAscan.py -in data1.fq.gz -in2 data2.fq.gz -alsgenescanner -out
/path/to/outdir
~~~

* ALSgeneScanner is part of the DNAscan analysis framework

Our table of sensitivity, specificity and accuracy (Table 2) means that the appropriate cut-off can be used to interrogate data, depending on whether the aim is the exclusion of potentially harmful variants, or the detection of definitely harmful variants.

ALSgeneScanner puts a powerful bioinformatics tool, able to exploit the potentialities of next generation sequencing data in the hands of patients, ALS researchers and clinicians.

